# Inferring neuronal ionic conductances from membrane potentials using CNNs

**DOI:** 10.1101/727974

**Authors:** Roy Ben-Shalom, Jan Balewski, Anand Siththaranjan, Vyassa Baratham, Henry Kyoung, Kyung Geun Kim, Kevin J. Bender, Kristofer E. Bouchard

## Abstract

The neuron is the fundamental unit of computation in the nervous system, and different neuron types produce different temporal patterns of voltage fluctuations in response to input currents. Understanding the mechanism of single neuron firing patterns requires accurate knowledge of the spatial densities of diverse ion channels along the membrane. However, direct measurements of these microscopic variables are difficult to obtain experimentally. Alternatively, one can attempt to infer those microscopic variables from the membrane potential (a mesoscopic variable), or features thereof, which are more experimentally tractable. One approach in this direction is to infer the ionic densities as parameters of a neuronal model. Traditionally this is done using a Multi-Objective Optimization (MOO) method to minimize the differences between features extracted from a simulated neuron’s membrane potential and the same features extracted from target data. Here, we use Convolutional Neural Networks (CNNs) to directly regress generative parameters (e.g., ionic conductances, membrane resistance, etc.,) from simulated time-varying membrane potentials in response to an input stimulus. We simulated diverse neuron models of increasing complexity (Izikivich: 4 parameters; Hodgkin-Huxley: 7 parameters; Mainen-Sejnowski: 10 parameters) with a large range of variation in the underlying parameter values. We show that hyperparameter optimized CNNs can accurately infer the values of generative variables for these neuron models, and that these results far surpass the previous state-of-the-art method (MOO). We discuss the benefits of optimizing the CNN architecture, improvements in accuracy with additional training data, and some observed limitations. Based on these results, we propose that CNNs may be able to infer the spatial distribution of diverse ionic densities from spatially resolved measurements of neuronal membrane potentials (e.g. voltage imaging).

## 1. Introduction

The brain analyzes information using many types of neurons with diverse firing properties, giving rise to the complex processing that occurs in the brain. Neuronal firing in response to a given input is in part dependent on the biophysical properties of the neuron, including the spatial distribution and density of ion channels along its membrane [1, 2]. To better understand the underlying mechanisms of neuronal activity, we need to model single neurons accurately, including the location and density of these ion channels. However, determining these parameters empirically is technically challenging, especially across complete neuronal arbors [3, 4, 5]. This poses a major challenge to simulate neurons using computational methods [6]. A common approach to overcome this challenge is to use neuronal models with ion channel densities described by adjustable parameters [7, 8, 9, 10]. These parameters can then be fit to experimental recordings, thus constraining the densities of the ion channels to reproduce features of experimentally observed membrane potentials [11]. These fitted parameters are predictions of the recorded neuron’s ion channel distributions, and can be used to implement a computational model of the neuron [7, 12, 13, 14].

Currently these types of models are fitted using evolutionary algorithm such as multi-objective optimizations (MOO) [8]. However, these optimization methods have their limitations. First, the results depend heavily on the metrics used to compare the model’s output to the neuronal recordings, with different metrics often producing inconsistent results [6]. Second, even when using the same metric, variations in initial conditions can lead to drastically different predictions for the ion channel distributions. Finally, MOO lacks a well-defined and differentiable objective function, and thus requires the use of *ad hoc* optimization schemes that are typically less efficient. CNNs has been used to infer cosmological parameters from simulations [15]. Here we explore a new approach where we use CNNs to directly predict the ion channel distributions from observed or simulated membrane potentials.

## 2. Methods

### 2.1. eFEL features from neuronal waveforms

To compare voltage waveform we used the electrophysiology Feature Extraction Library (eFEL) [16, 17]. We extracted the following features from each waveform: mean frequency of action potentials (APs), adaptation index, time to first AP, mean AP amplitude, average hyperpolarization depth, and AP half width. These features were chosen similarly to those used in here [8]. The eFEL features have been used in this work for two purposes: providing loss for MOO convergence (see below) and to evaluate the quality of CNN predictions.

### 2.2. Workflow for training and validating the CNNs

The CNNs used in our method are trained on data generated by computational simulations of neuronal activity. Our simulations take two inputs: the parameters defining the biophysical properties of the neuron (ion channel densities), and a stimulus to be applied to the neuron during whole-cell current-clamp recordings. The results presented here were obtained using the chirp-type stimulus shown in Fig A1 in the appendix.

The resulting membrane potentials paired with the input neuronal parameters formed the CNN training set. The overview of the workflow steps used in this paper are summarized in Fig. 1. First, up to 2 million examples per neuronal model were generated by uniformly randomly sampling a multidimensional parameter space, and the generated examples were split into training/validation/test sets. The training and validation examples were then used to train 32 independent CNN models, each of which had the same architecture but started with a different initialization of the weights and processed the training data in a different order. The 32 CNN predictions made for the ‘test’ dataset were averaged, resulting in a single prediction of the parameter values, drawn as a green banner with dashed outline in Fig. 1. The CNN-predicted parameters were compared using the ground truth values and the residual errors averaged over the sampled parameter space. We have also compared the features of the voltage waveform using the eFEL [17] (see Section 2.1) between the waveforms obtained from simulation using the CNN-predicted parameters, and those obtained using the ground-truth parameters.

**Fig. 1:**
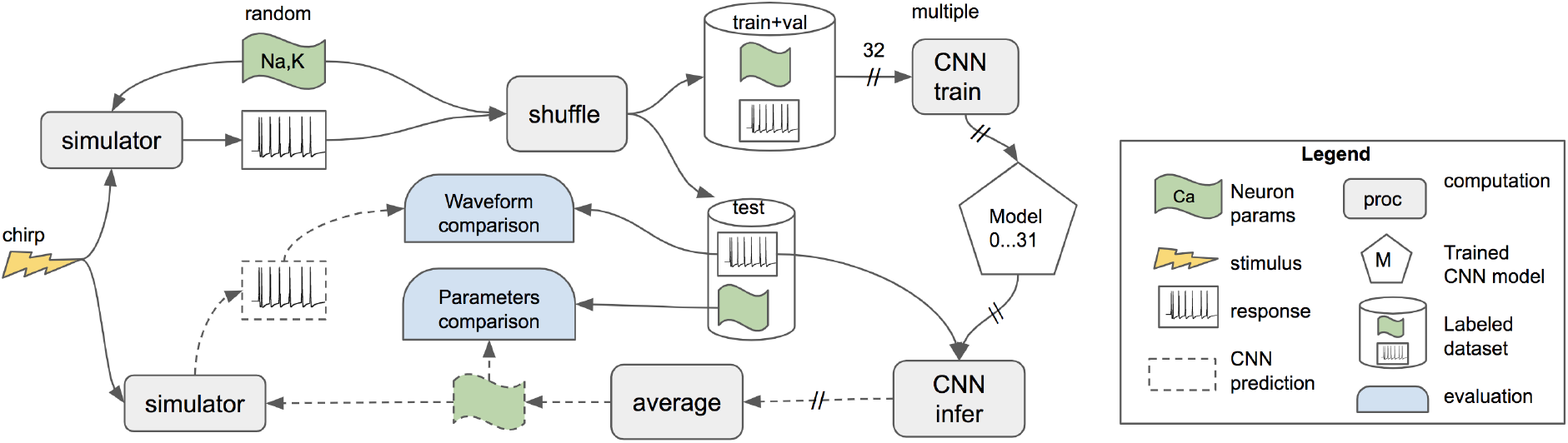
Workflow diagram describing generation of voltage traces, CNN training, prediction of ion channel densities from voltage traces input, and validation of CNN predictions.

### 2.3. Neuronal models

In this paper, we evaluate CNN-based methods using three neuronal models: (i) the simple spiking neuron Izhikevich model [18] which has four parameters (Table A3); (ii) Hodgkin and Huxley [19] dispersed channels model assuming a ball and stick morphology (Tables A4, A5); (iii) a complex Mainen-Sejnowski model [20] with realistic pyramidal cell morphology and some additional passive conductance e.g. *g_l_* (Table A6). We control the complexity of each neuronal model by choosing only a subset of the available free parameters to vary, with the rest held at fixed values or latched to another parameter’s value (for example, we chose to use the same leak conductance at the soma and the dendrite of the Hodgkin-Huxley ball and stick model). The additional description of the neuronal models can be found in the appendix data. We evaluated a total of 8 neuronal models, of which only 3 are described here. The other 5 variants are described in the appendix.

### 2.4. Generating training datasets

We generated somatic membrane potentials using the NEURON simulator [21]. The names and ranges of our three models’ free parameters are described in the Tables A3–A6. To invoke a voltage response we used the chirp-type stimulus - Fig. A1. The simulated waveforms were accepted for the CNN-training if there were (i) no spikes during the hyperpolarization phase of the stimulus, and (ii) less than 30 total spikes during the chirp phase of the stimulus. Note that, in the CNN training process, we disregarded the cell’s response during the initial hyperpolarizing current.

The number of examples used to train the CNNs varied from 1 × 10^5^ examples for the Izhikevich 4 parameter dataset to 1.5 × 10^6^ examples for the Mainen and Sejnowski 10 parameter dataset (see Tables A3–A6 for the details). The dynamic ranges of the physical values for each varied parameter were chosen to be approximately between 0.5 times the base value or 2 times the base value used by the authors of the respective neuron models [18, 20]. Some exception were made when the same ion conductances was varied in neighbouring compartments (e.g. *g_Na_dend_* and *g_Na_soma_* in Table A6). The non-varied parameters were either kept constant at the base values or co-varied with one of the varied parameters as described in Section 2.3.

### 2.5. CNN model

A machine learning regressing network is a function *F*(*X*(*t*)|***θ***) = **Y**, where *X*(*t*) is the temporal pattern of voltage fluctuation of a biological neuron membrane, called a “trace” or “waveform” for short. *F* also depends implicitly on the CNN weights ***θ*** which are determined during the training process. The output **Y** is a multi-valued floating point vector corresponding to the predicted parameter values. Since the activation of the last fully connected (FC) layer was set to be 1.2 × tanh the range of accessible **Y**s was [−1.2,1.2].

Convolutional Neuron Networks (CNN) have been successfully used in a variety of ML-tasks, e.g., 2D image processing. Since a biological neuron’s activity (a trace) is typically recorded as a single valued time series, the CNN was constructed from several 1-dimensional (1D) CNN layers interleaved with 1D-Pooling layers to extract information about repetitive features (neuron spikes). The following block of FC layers was intended to capture the global features of the trace. The last FC layer returned regressed parameters (e.g., ionic conductances, membrane resistances, etc.,) modelling a neuron. Those physical parameters were linearly scaled to the unit-less range [−1,1]. The predictor *F*(.) had the capacity to produce values with amplitudes up to 20% larger than necessary by design, to allow out-of-range predictions.

The natural amplitude of the membrane potential varied between [−80, 50] mV. Since we expect the nonspiking portion of a trace to contain more information about ionic channels, we shifted each trace so that its minimum value was +1mV and took the logarithm, as follows:

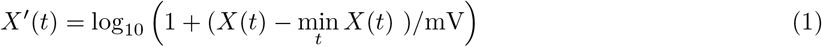

This guarantees that the range of input values *X*′ is limited to [0,3] and only a small fraction of this range is designated to neuron spikes. Also, it reduces sensitivity to the drift of holding current, which is expected to vary between stimulations. The dimension of *X*(*t*) was 9000 time bins.

The {*X*(*t*),**Y**} dataset was randomly divided into train/validate/test sets in a 7/1/1 ratio. During the CNN training, we reduce the learning rate at each step if there was no improvement in the validation loss, and terminate learning after no further improvement of loss was observed. We executed 32 independent trainings on the same train and validation datasets, each time differently shuffling the examples. The final CNN-predictions were averaged over 32 sets of learned weights ***θ***.

The optimal configuration of the layers of the CNN and the training method were determined by means of a random search of the hyper-parameter space [22]. We executed 780 CNN-training experiments with 14 randomly varied hyper-parameters (hyper-parameters include number of CNN and FC layers, number of features per layer, optimizer type, loss function, etc.; see the appendix for more details). Each training lasted between 4 and 6 GPU-hours, depending on how quickly the loss stabilized. The best model was not necessarily the one with the smallest one-time training loss value. By repeatedly training the same model using random permutations of the same data, we confirmed that training was stable, and that the final validation loss achieved by each training instance was minimally variable across permutations.

The best-performing CNN model we tested has 744,000 weights (***θ***) and consists of 6 CNN, 3 Pool, and 7 FC layers. The CNN layers had 18 or 25 features, the flatten layer had 45,000 units, the feature count for the FC layers varied between 148 and 46 (with deeper layers having fewer features), the last FC layer output had the dimension equal to the number of regressed parameters (**Y**). CNN kernel size was 5, pool size was 3, the dropout rate between FC layers was of 0.01, the optimizer was ADAM [23], the learning rate reduction factor was of 0.14, and mean absolute error (MAE) loss was used.

### 2.6. Multi-Objective Optimization (MOO)

MOO, as implemented in BluePyOpt [16], is the standard method for fitting neuronal models to *in vitro* recorded data. This evolution based method constrains the free parameters of the neuronal models so it’s output will have similar objective as the target neuronal data. This is done by minimizing the RMS between features (eFEL) [17] of the voltage waveform - the objectives of the optimization. BluePyOpt’s implementation of MOO attempts to minimize each objective using pareto-optimality conditions as described here [24]. When the optimization terminates, a group of solutions (neuronal model parameters predictions) with high similarity to the target data are suggested. Here the target neuronal data is first generated using the base values of the relevant neuronal model (Tables A3,A4, A6).

### 2.7. Evaluation of CNN and MOO accuracy with eFEL

eFEL features were also used to evaluate and compare the quality of CNN and MOO parameter inference. For each sample in the test set, we computed the eFEL features listed in Sect. 2.1 from the ground-truth waveform as well as from the CNN-predicted waveform (see Fig. 1). The mean (*μ*) and standard deviation (*σ*) of the differences between eFEL features from the ground-truth and CNN-predicted waveforms across the whole test set were estimated for each feature. The total error *ϵ* of CNN or MOO prediction was computed by adding both moments in quadrature, assuming they represent two statistically independent measures of the error of predictors.

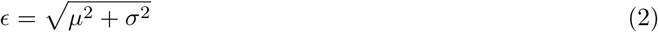

We note that the eFEL library was unable to compute some of the chosen features for some waveforms. In such cases we would discard the input. This affected 25%, 10%, and below 1%, of the Izhikevich, Hodgkin-Huxley, and Mainen test datasets, respectively. To establish a baseline, 50,000 waveforms produced by each neuron simulator with randomly varied parameters for the purpose of CNN training has been set aside, divided in to pairs and eFEL feature difference was computed for each pair. These distributions were further refined by eliminating outlying comparison values that exceeded the 10 standard deviation interval. Finally, the mean and standard deviation was recomputed for the filtered distributions of feature value differences. The baseline error (eq. 2) was computed for each dataset.

## 3. Results

### 3.1. CNNs can accurately predict ion channel densities of neuronal models

We trained the same CNN-model on data from each of 8 different neuronal models. Here we will discuss the results for three selected models, in order of increasing model complexity. The CNN predictions are made for ‘test’ examples not used in the CNN-training process.

#### 3.1.1. Izhikevich - phenomenological spiking neuron model

We started with the simplest model, the Izhikevich neuron. Izhikevich is a spiking neuron model described by 2 differential equations [18] and does not account for morphological effects; i.e. the Izhikevich model is a point cell model. The Izhikevich model can be used to represent a wide range of firing patterns by varying the parameter values. Spiking neuron models such as Izhikevich are mostly used to represent neurons in a large scale neuronal simulation with many point neurons. The Izhikevich model is not readily interpretable in terms of biophysical properties of the simulated neuron.

The predictions for the Izhikevich dataset are shown in Fig. 2a). The regression residues, defined as the difference between the truth and predicted value, have a RMS between 1-2%, averaged over the whole sampled dynamic range of respective parameters [−1,1]. Despite training on only 3 × 10^4^ examples, the CNN achieved high prediction accuracy. This can be attributed to the relative simplicity of the Izhikevich model, where the parameters contribute directly to the voltage response. After training, we used the predicted values with either high or low error to generate voltage responses and overlay them with the ground truth voltage response (Fig. 4a).

**Fig. 2:**
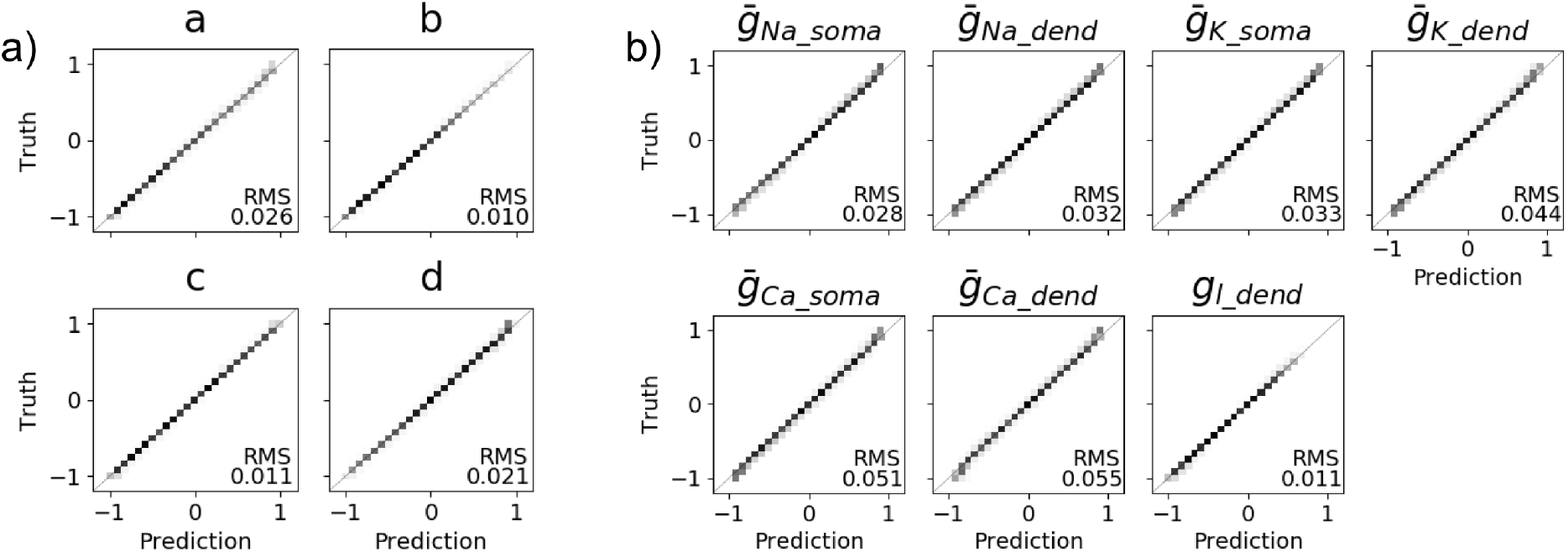
Correlation between neuron parameters used by the simulator (ground truth) and the values predicted by the CNN-model trained on the respective dataset. The average RMSE of the difference between the truth and predictions is shown in the lower right corner of each parameter plot. a) Izhikevich simulator with 4 varied parameters, b) Hodgkin-Huxley simulator with 7 varied parameters.

#### 3.1.2. Hodgkin-Huxley ball and stick compartmental model

Next, we simulated a reduced neuronal morphology: a sphere connected to a cylinder, representing a simplified soma and dendrite, respectively. The adjustable parameters in the Hodgkin-Huxley 7-parameter model were the maximal ionic conductances of various ion channels in each of the two compartments. This model was used to study whether a CNN could be trained to separate contributions from the same ion channels in two different compartments. We trained the CNN with 4.5 × 10^5^ examples. The CNN predictions of the parameter values are shown in Figs. 2b, 4b. The CNN trained for the Hodgkin-Huxley 7-parameter model made accurate predictions with small residues below 6%. The highest prediction accuracy (residue RMS of 1.1%) was achieved for the leak conductance at the dendrite *g_l_dend_*. This is the only conductance in the Hodgkin-Huxley 7 parameter model that was was not allowed to take different values at the soma and the dendrite. The most difficult parameters to predict were the correlated somatic and dendritic calcium conductances (residue RMS of 5.1% and 5.5%, respectively). We attribute this to the challenge of distinguishing the contribution of the calcium conductance in dendritic compartments to the voltage response recorded at the soma.

To further study the CNN’s ability to separate the contributions of conductances originating from different compartments, we used two Hodgkin-Huxley ball and stick models with only 4 varied parameters. In the first of these 4 parameter models (Hodgkin-Huxley 4parE), each parameter controlled the conductance of a different ion channel type. In the second (Hodgkin-Huxley 4parH), two ionic conductances (Na, K) were varied separately at each of two different compartments. Although the Hodgkin-Huxley 4parH predictions were more closely correlated with the true values than the 4parE predictions (see Fig. A2b, both models’ parameters were predicted with high accuracy. Next, we wanted to check the limits of the CNN trained model with Hodgkin-Huxley ball and three sticks morphology. We added another two compartments to the morphology to simulate a basal dendritic arbor with different physical dimensions and topology. In this model, we varied 10 parameters (Hodgkin-Huxley 10par) which describe the conductances at the 3 different compartments (see Fig.A3 and Table A5). Our CNN model was able to predict only 7 parameters with residues below 10%, despite training on 1.4 × 10^6^ examples for over 10 GPU-hours. It seems we reached the limit of CNN predictions when using only one stimulus and voltage response only from the soma.

#### 3.1.3. Mainen and Sejnowski biophysically detailed compartmental model

Finally, we studied how the trained CNN would predict the ionic conductances of a biophysically detailed model with realistic morphology such as Mainen and Sejnowski [20]. We tested this neuronal model with gradually increasing complexity by varying 4, 7, and 10 parameters (Table A6, Figs. A4, A5, and 3, respectively). When 4 parameters were varied, prediction errors were less than 4%. Increases in the average prediction error were seen as more parameters were varied (Table A6). The 10 parameter model introduced the new challenge of multiple variables controlling the same ionic conductance in *the same compartment*. For example, the total potassium conductance can be controlled by the following 3 conductances in several different combinations: calcium, potassium M-current, and calcium-activated potassium channels (*KCa*). This redundancy results in a fundamental lack of parameter identifiablity. Indeed, these three parameters proved to be the most strenuous to resolve: the CNN was able to determine them with residues of 10%, 29%, and 13%, respectively. One of the advantages of using CNNs is that we can easily analyze the correlations between the predicted parameters to gain insight into their interdependence (Fig. 3b). Despite the relatively high errors in the Mainen 10 parameter predictions, the produced waveforms were qualitatively similar (Fig. 4c). However, since the Mainen model is sufficiently complex to represent many of the underlying mechanisms of neuronal firing, this is an example of how CNN can be a useful tool to predict ion-channels densities of neurons from *in vitro* voltage responses.

**Fig. 3:**
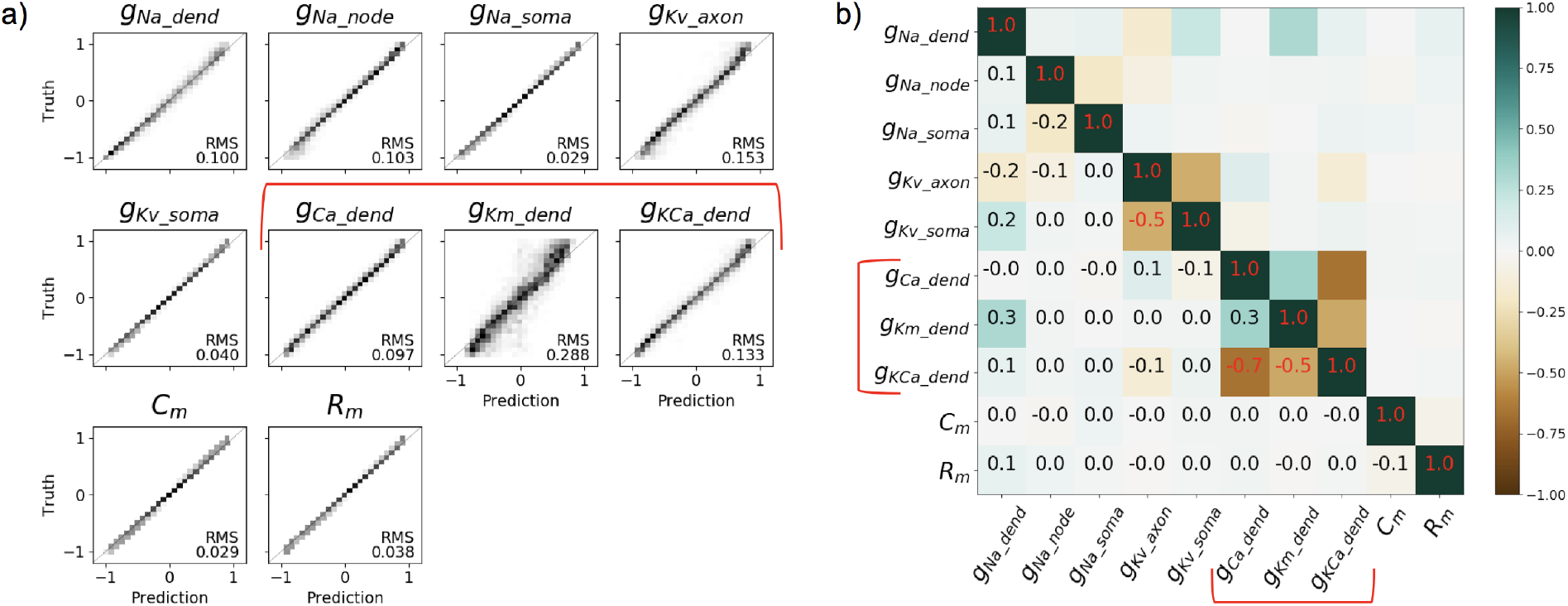
CNN-predictions for 10-parameter Mainen dataset. (a) The correlation between the truth and predictions. (b) correlation between residua of predictions depicted by color scale, where 0 means no correlation and (-)1 means strong (anti)correlation. The brackets mark most difficult to predict, highly correlated triplet of dendrite conductances: Ca, K_*m*_, and KCa.

**Fig. 4:**
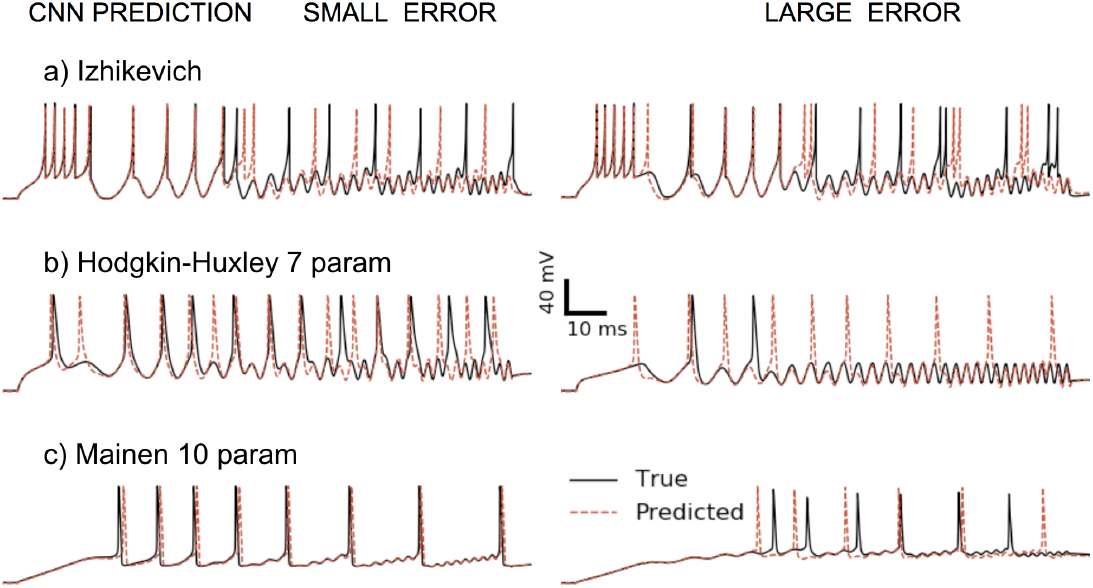
Examples of voltage traces generated from CNN predictions with varying errors, overlaid with the true voltage. Small errors are 0.5 STD below the populations mean.

### 3.2. CNN outperforms Multi Objective Optimization

We tested if the CNN predictions for the free parameters of neuronal models could be an alternative to MOO - the standard method to fit model parameters to voltage responses. Here our optimization used six eFEL features similar to the ones used in here [8]. We ran MOO for each of the 3 models discussed above, terminating each optimization after 10 generations with no significant improvement, or when the number of parameter sets evaluated during the optimization process exceeded the number of samples used to train the CNN on the same neuronal model (Tables A3, A4 and A6). The target voltage responses were the ones obtained with the base parameter values, shown in the same Tables. After the optimization terminated we compared eFEL features between the target trace and the traces generated by the 32 best parameter predictions found during MOO. We also made the same comparison for the 32 predictions made by the 32 trained ML models on the sample whose parameters were closest to the base values in (Tables A3, A4 and A6). The predicted parameters from both MOO and the CNNs produced traces with eFEL features similar to the respective target traces are shown in Fig. 5.

**Fig. 5:**
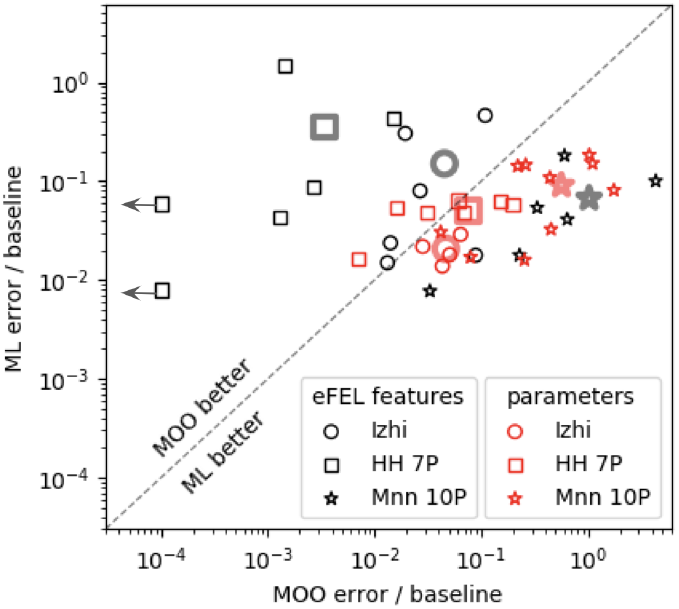
Comparison of relative error of MOO vs. CNN predictions. The larger fainter symbols denote average over respective observables.

To qualitatively compare the performance of MOO and the CNN, we used two criteria discussed in methods: similarity of the predicted waveform to the target data and the distance of prediction in the parameter space. In the features comparisons MOO did better only for the simple models (black symbols in Fig. 5). In the parameter loss criteria, overall, CNN outperformed MOO in all models (red symbols in Fig. 5). We note that in the Hodgkin-Huxley 7-paramater model, the CNN predictions did a poor job of reproducing the eFEL features of the target trace in comparison to MOO, despite the fact that the parameters were determined to slightly better accuracy than MOO. In summary, we conclude that (i) in simpler models, MOO performs better than our CNN in fitting features, (ii) our CNN does better in fitting the parameters on average in all models, (iii) for very complex models, a strategy based on accurately fitting the parameters is more effective than one that optimizes for reproduction of eFEL features of voltage waveforms.

## 4. Discussion

### 4.1. CNNs can accurately predict ion-channels densities of neuronal models

In this study we explored applicability of CNN to predict ionic channels’ conductances of neuronal models from their voltage response. We trained the same CNN on datasets from 3 different neuronal models: spiking neuronal models, Hodgkin-Huxley reduced morphology, and biophysically detailed models. The CNN with optimized hyper-parameters was able to recover majority of parameters for all of the 8 models we trained it on with varying accuracy. The most challenging to predict were parameters that were highly correlated, either due to having the same conductances in several compartments or several conductances at the same compartment that control the same ion flux. This issue was more prevalent as the complexity of the models increased. For the two neuronal models with 10 parameters (Figs. 3 and A3). CNN predicted several parameters with large residues of 30%-50%, indicating this might be the limit of neuron complexity when using only one stimulus and recording only one voltage at the soma. We verified the waveforms generated from the CNN predictions with small or large prediction errors (Fig. 4) and qualitatively the earlier seemed to better agree with the ground truth. Furthermore, we computed the features of the waveform using the eFEL package (see methods) to quantitatively compare between the waveforms generated from the CNN predictions (Fig. 5).

### 4.2. CNNs outperforms the current standard method in predicting ion-channels densities

The main challenge in fitting neuronal models to the waveforms is that multiple sets of parameters may lead to the similar neuronal output [25, 6], which is caused by model non-linearity and/or correlation between the parameters. We are aiming to find the solutions a recorded neuron is using. Currently, the most common way to fit models parameters to *in vitro* neuronal recordings is using MOO as initially described here [8] and later standardized in BluePyOpt [16]. This method is used either for constructing large scale neuronal simulations [9] or to represent a recorded neuron as a compartmental model [7]. The MOO optimization is computationally intensive and needs to be repeated for every new target waveform. On average, CNN predictions for neuronal model’s parameters were more accurate than MOO for all the models tested in this study (3 examples in Fig 5). For simple models both MOO and CNN had comparable small prediction errors. For more complex models (Mainen 10 parameters), MOO found a different solution that generated similar waveforms to the target data but resulted in large parameter errors.

In addition to accuracy, there are other advantages of using CNNs over MOO: first, a trained CNN can predict neuronal parameters for any target waveform in a fraction of a second while MOO method requires to run a costly optimization for each new target data, often for many CPU-days. Second, MOO convergence highly depends on the choice of the objectives (features of the waveform) and it is not clear how to choose the right ones out of more than 200 offered by the eFEL package. Third, CNN predictions applied to a large dataset provide information about statistical error of predictions and correlations between parameters (Fig. 3a,b). It provides new insight in neuron models, helps in improving them, and allows quantitative comparison of accuracy of different models.

### 4.3. Future directions and applications

To the best of our knowledge, this is the first demonstration of applying CNNs to predict parameters of neuronal models. There are several improvements that can be applied to our method to solve even more complex and biophysically detailed models: i) Adding more stimulation to generate a more versatile response from the neuronal simulator [26], ii) recording from additional locations on the neuron, either using the cell attached configuration [27] or direct voltage imaging of the entire cell, iii) reducing the dimensionality by only predicting sub-groups of the varied parameters and pharmacologically blocking other conductances as described here [28]. Moreover, we would like to generalize CNN-based predictor to also indicate the level of mismatch between the input waveform and the CNN training dataset. In the future we want to apply the method describe here to spatially map the ion channels of recorded neurons to better understand the underlying mechanisms of their firing patterns.

## 5. Acknowledgements

This research was supported in part by SFARI grant 513133 and NIH MH112117, and by previous breakthroughs obtained through the Laboratory Directed Research and Development Program of Lawrence Berkeley National Laboratory under U.S. Department of Energy Contract No. DE-AC02-05CH11231. This research used resources of the National Energy Research Scientific Computing Center (NERSC), a U.S. Department of Energy Office of Science User Facility operated under Contract No. DE-AC02-05CH11231.

## Appendix A. Supplementary Methods

### A.1. Stimulus

The chirp stimulus used across this study is a sinusoidal wave with continuously varying frequency and amplitude:

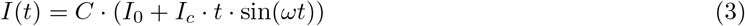

where *I_c_* = 11 nA/s, *ω* = 0.64 rad/ms, *I*_0_ = 0.5 nA, *I_c_* was adjusted to control the amplitude for each neuronal model so it would generate APs. We added a hyperpolarizating current lasting 110 ms to prevent spontaneous spikes prior to the onset of the stimulating chirp, as well as an 80 ms period of no current injection post-chirp to allow for some spontaneous (i.e., non stimulus-driven) activity. The whole current injection is shown in Fig. A1.

**Fig. A1:**
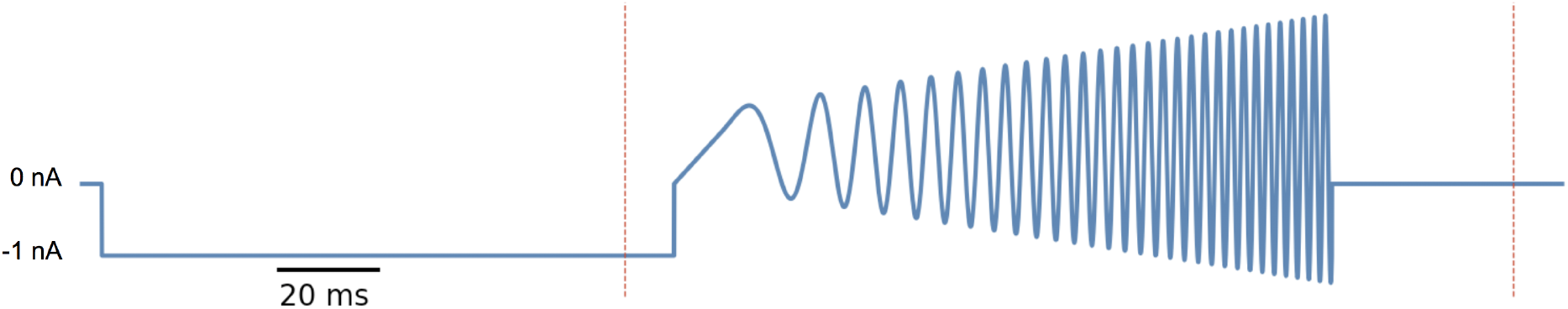
Stimulus used in our neuronal simulations. Red lines mark the range used for CNN training.

### A.2. Neuronal Models

All simulations were run in the NEURON simulation environment [21] version 7.5. All cell and ion channel models were obtained from ModelDB [29].

The Izhikevich model [18] was implemented as a `POINT PROCESS` in a zero-length cellular compartment in the NEURON simulator.

The Hodgkin-Huxley simulations used ion channel models from Mainen and Sejnowski, [20]: Na^+^, Ca^2+^, and fast Potassium (K_V_) channels with kinetic parameters held at their predefined values from the Mainen+Sejnowski cells (only the max conductances of each channel type were left as adjustable parameters). For the 5 parameter Hodgkin-Huxley cell, these channels were placed into a simple morphology with the soma represented by a compartment with 21 *μ*m length and diameter, and a dendrite of length 210 *μ*m and diameter 2.1 *μ*m. For the 10 parameter Hodgkin-Huxley cell, there was an additional basal dendritic structure consisting of two compartments, each with length 52.5 μm and diameter 2.1 *μ*m, both attached to the soma opposite the apical dendrite.

### A.3. CNN-prediction accuracy vs. training size

We used Mainen 10 parameter dataset for study of accuracy of ML-prediction as function of the size of training data set. The same ‘test’ data segment was set aside from the trainings and used as common reference for the inference. For each choice of the training data size we run 8 independent ML-trainings, on different subset of examples, shuffled differently. Only for the largest data size choice we just shuffled the examples. The results reported in table A1 are averaged over 8 trainings.

**Table A1:**
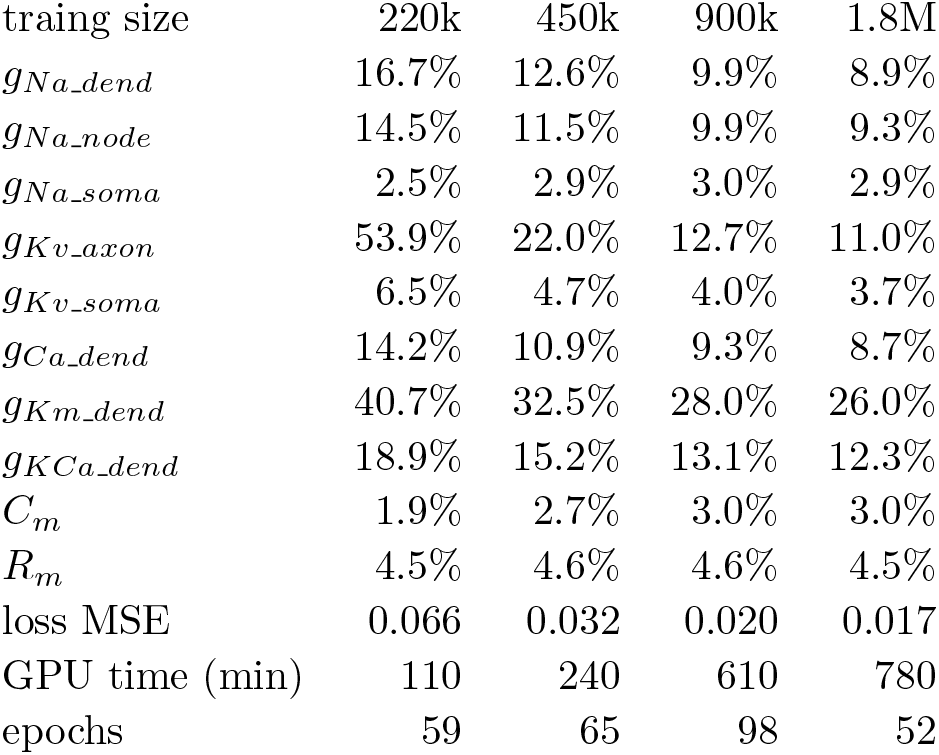
Accuracy of ML-prediction for Mainen 10param dataset as function of the size of the training data set.

### A.4. Optimization of CNN

The initial CNN-model used to develop the CNN code-base was defined by the following set of hypar-parameters (ver 1)

The CNN-model was constructed based on hyper-parameters as follows:

#### Listing 1

Construction of CNN model from hyper-parameters (pseudo-code)

~~~
h = Input(shape = (9000,1))
#….. *CNN–1D layers*
cnn_ker=hpar[‘conv_kernel’]
**for** dim **in** hpar [‘conv_filter’]:
    **for** j **in range**(hpar[‘conv_repeat’]):
        h= Conv1D (dim, cnn_ker, activation=‘linear ’, padding= ‘valid’)(h)
        h =LeakyReLU ()(h)
    h= MaxPool1D (pool_size=hpar [‘pool_len’])(h)
h=Flatten ()(h)
h = Dropout (hpar[‘dropFrac’])(h)
#…. *FC layers*
**for** i, dim **in enumerate**(hpar [‘fc_dims’]):
    h = Dense (dim, activation=‘linear’)(h)
    h =LeakyReLU () (h)
    h = Dropout (hpar[‘dropFrac’])(h)
#… *Output*
y= Dense (out_dim, activation=hpar [‘lastAct’]) (h)
y = Lambda(**lambda** val: val * hpar [‘outAmpl’]) (y)
~~~

**Table A2:** hypar-parameters for CNN model ver1 and ver693. ‘*’ - not varied

|  | Parameter | ver 1 | ver 693 |
| --- | --- | --- | --- |
| CNN params | conv_filter | [6,12,18,24,30,36] | [18, 25] |
|  | conv_kernel | 5 | 5 |
|  | conv_repeat | 2 | 3 |
|  | pool_len | 3 | 7 |
| FC params | fc_dims | [20, 10, 10] | [148, 131, 114, 97, 80, 63, 46] |
|  | lastAct* | tanh | tanh |
|  | outAmpl* | 1.2 | 1.2 |
|  | dropFrac | 0.02 | 0.01 |
| Training | lossName | mse | mae |
|  | optimizerName | adam | adam |
|  | reduceLR_factor | 0.3 | 0.14 |

### A.5. Supplementary figures and tables

**Table A3:**
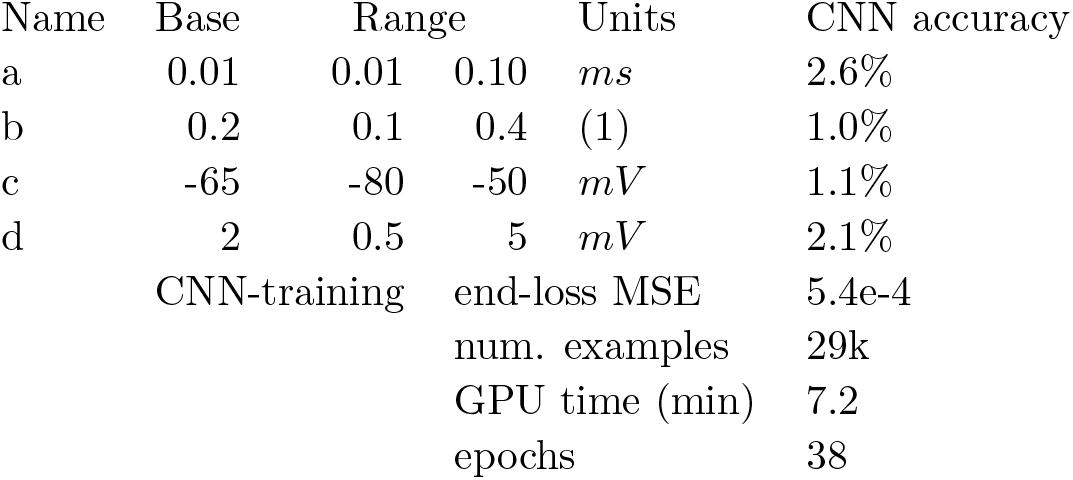
Ranges of parameters used by Izhikevich model and accuracy of CNN predictions, averaged over 32 trained models.

**Table A4:**
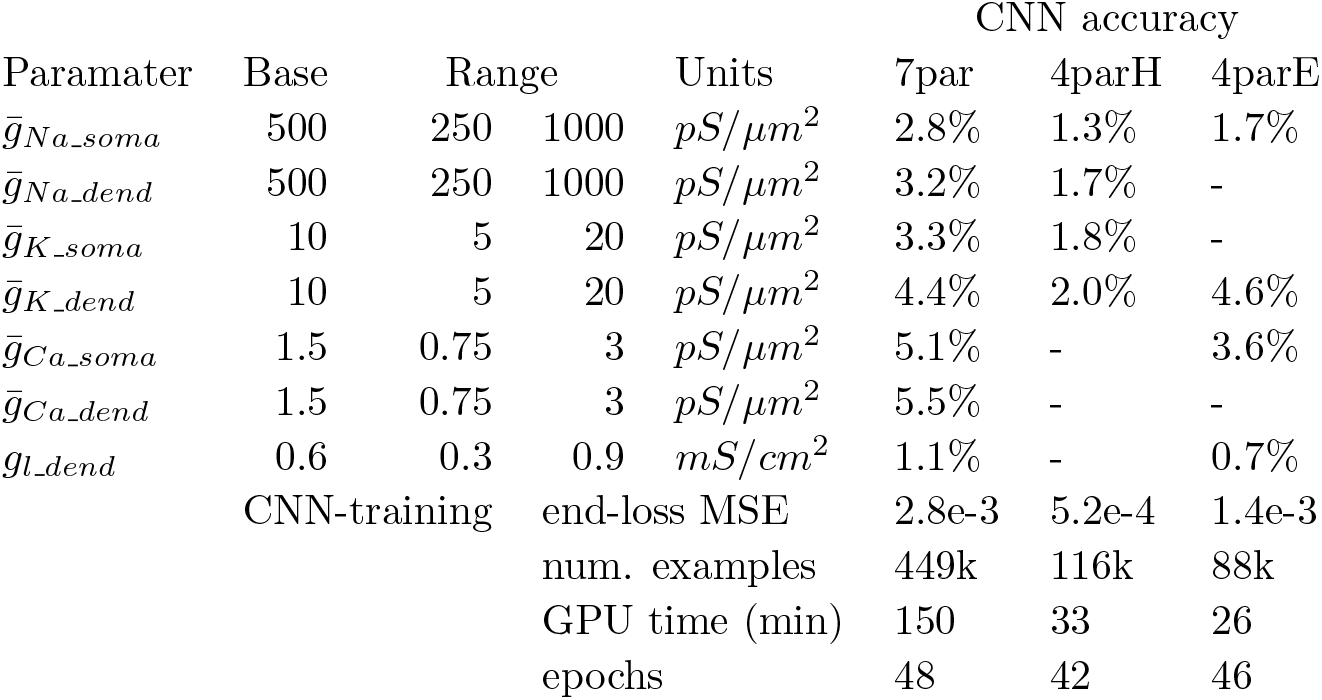
Ranges of 4 or 7 parameter sets used by Hogkins-Huxley ball+stick model and respective CNN prediction accuracies. The value of *g_l_soma_* was locked to *g_l_dend_*.

**Table A5:**
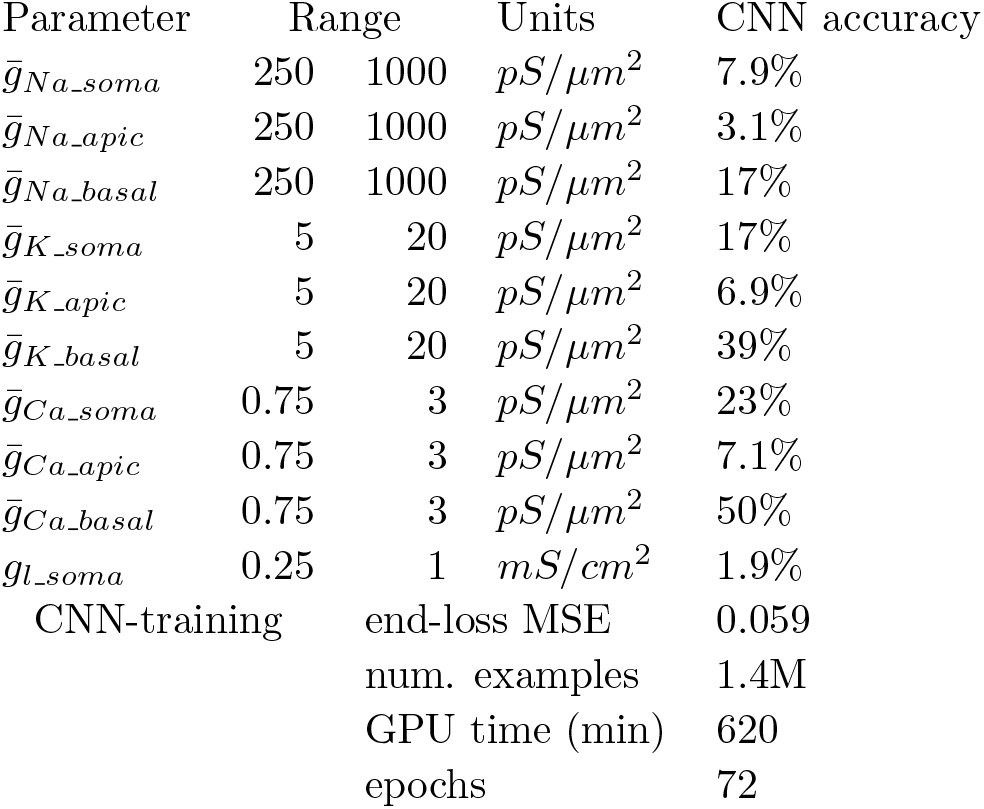
Ranges of 10 parameters used by Hodgkin-Huxley two dendrites model and respective CNN prediction accuracies. The value of *g_l_apic_* and *g_l_basal_* were locked to *g_l_soma_*.

**Fig. A2:**
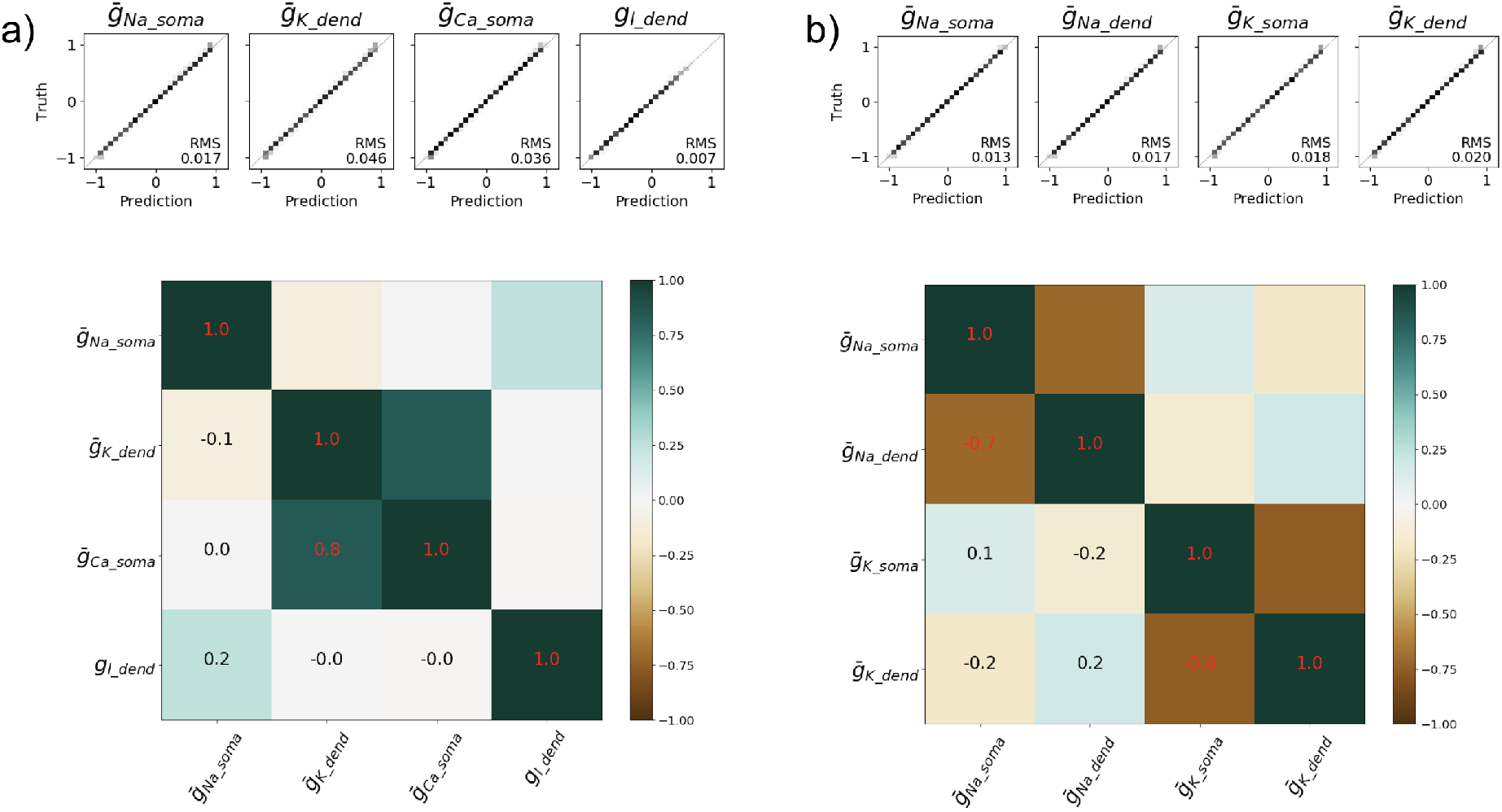
Comparisons of CNN predictions for Hodgkin-Huxley 4parE (a) and 4parH (b) with 2 different subsets of varied parameters described in Table A4

**Fig. A3:**
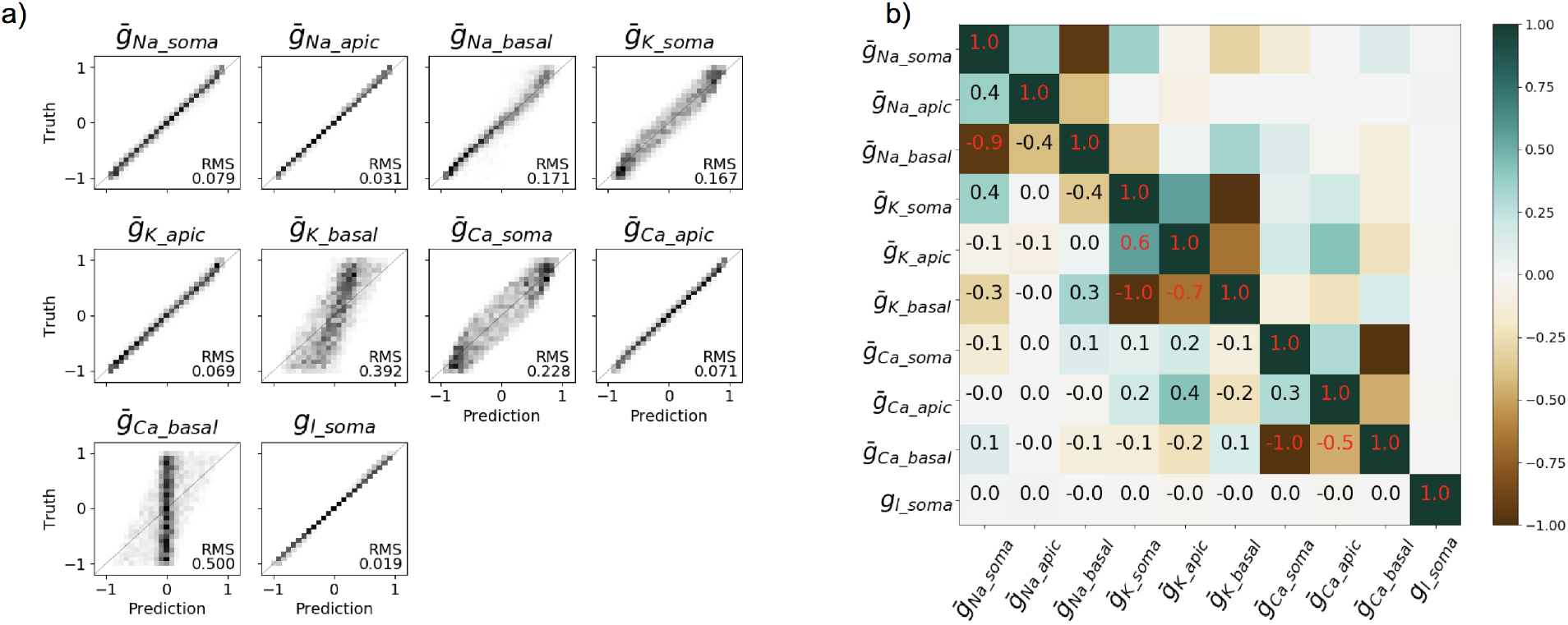
CNN prediction of 10 parameters for HodgkinHuxley ‘ball with 2 sticks’ geometry was only partially successful. The low sensitivity to 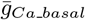 (left panel) is most likely cause by 90 % anti-correlation with 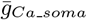 (right panel).

**Fig. A4:**
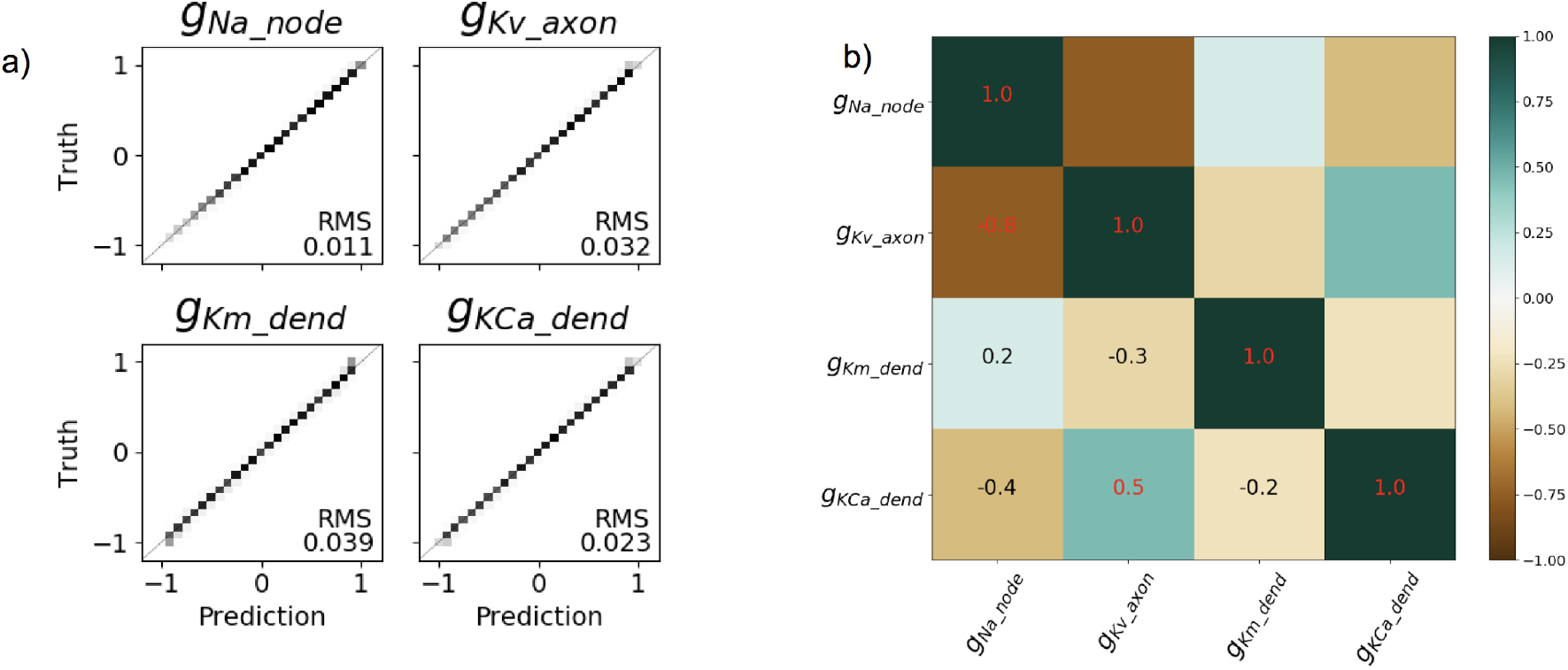
CNN prediction of Mainen 4-parameter neuron model.

**Fig. A5:**
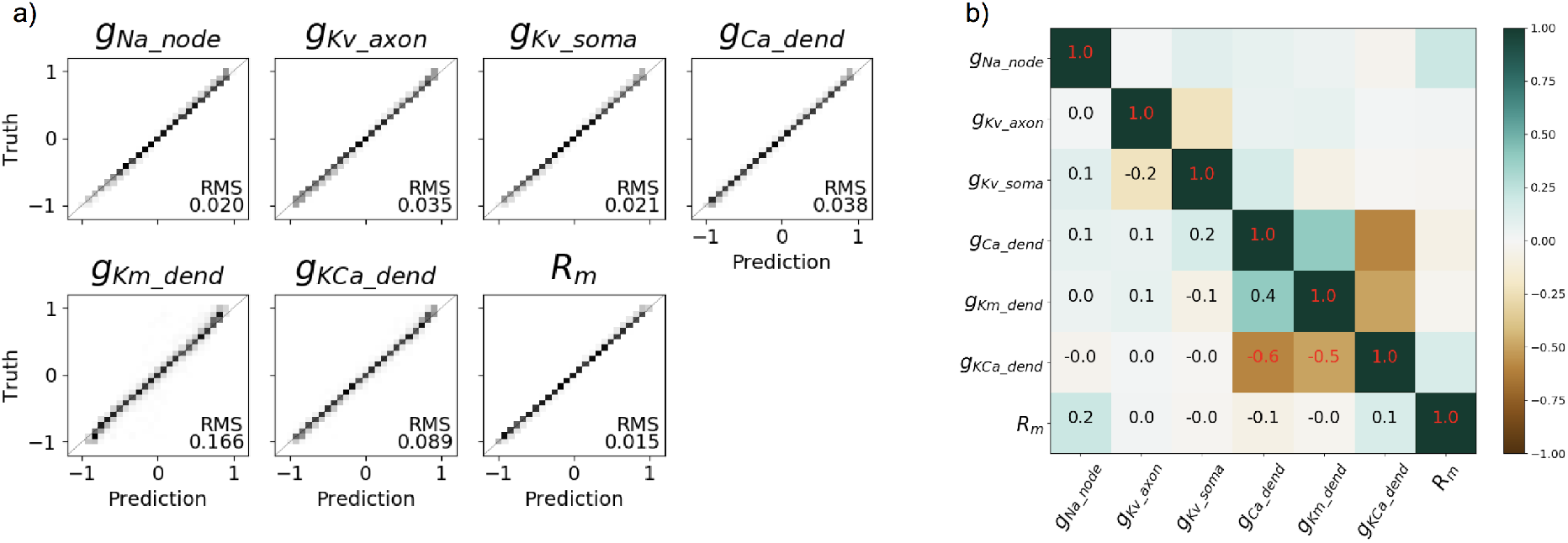
CNN prediction of Mainen 7-parameter neuron model.

**Table A6:** Ranges of 4,7 or 10 parameter sets used by Mainen neuron simulator and respective CNN prediction accuracies.

| Paramater | Base | Range |  | Units | CNN accuracy |  |  |
| --- | --- | --- | --- | --- | --- | --- | --- |
|  |  |  |  |  | 10par | 7par | 4 par |
| $gNa_{dend}$ | 20 | 10 | 250 | $S/cm^2$ | 10% | - | - |
| $gNa_{node}$ | 30 | 15 | 80 | $kS/cm^2$ | 10% | 2.0% | 1.1% |
| $gNa_{soma}$ | 0.02 | 0.01 | 60 | $kS/cm^2$ | 2.9% | - | - |
| $gKv_{axon}$ | 2 | 1 | 5 | $kS/cm^2$ | 15% | 3.5% | 3.2% |
| $gKv_{soma}$ | 0.2 | 0.1 | 2 | $kS/cm^2$ | 4.0% | 2.1% | - |
| $gCa_{dend}$ | 300 | 150 | 600 | $mS/cm^2$ | 9.7% | 3.8% | |
| $gKm_{dend}$ | 100 | 50 | 200 | $mS/cm^2$ | 29% | 17% | 3.9% |
| $gKCa_{dend}$ | 3 | 1.5 | 6 | $S/cm^2$ | 13% | 8.9% | 2.3% |
| $C_m$ | 0.6 | 0.3 | 1.5 | $\mu F/cm^2$ | 2.9% | - | - |
| $R_m$ | 30 | 15 | 60 | $k\Omega cm^2$ | 3.8% | 1.5% | - |
| CNN-training |  |  |  | end-loss MSE | 0.023 | 6.6e-3 | 9e-4 |
|  |  |  |  | num. examples | 780k | 400k | 77k |
|  |  |  |  | GPU time (min) | 310 | 110 | 20 |
|  |  |  |  | epochs | 51 | 48 | 38 |

